# Spyglass: a framework for reproducible and shareable neuroscience research

**DOI:** 10.1101/2024.01.25.577295

**Authors:** Kyu Hyun Lee, Eric L. Denovellis, Ryan Ly, Jeremy Magland, Jeff Soules, Alison E. Comrie, Daniel P. Gramling, Jennifer A. Guidera, Rhino Nevers, Philip Adenekan, Chris Brozdowski, Samuel R. Bray, Emily Monroe, Ji Hyun Bak, Michael E. Coulter, Xulu Sun, Emrey Broyles, Donghoon Shin, Sharon Chiang, Cristofer Holobetz, Andrew Tritt, Oliver Rübel, Thinh Nguyen, Dimitri Yatsenko, Joshua Chu, Caleb Kemere, Samuel Garcia, Alessio Buccino, Loren M. Frank

## Abstract

Scientific progress depends on reliable and reproducible results. Progress can be accelerated when data are shared and re-analyzed to address new questions. Current approaches to storing and analyzing neural data involve bespoke formats and software that make replication and reuse of data difficult. To address these challenges, we created Spyglass, an open-source data management and analysis framework written in Python. Spyglass provides reproducible pipelines for common neuroscience analyses and sharing of raw data, intermediate analyses, and final results within and across labs. Spyglass uses the Neurodata Without Borders (NWB) standard and includes pipelines for spectral filtering, spike sorting, pose tracking, and neural decoding. Spyglass can be extended to apply existing and newly developed pipelines to datasets from multiple sources. We demonstrate these features in the context of a cross-laboratory replication by applying advanced state space decoding algorithms to publicly available data.

New users can try out Spyglass on a Jupyter Hub hosted by HHMI and 2i2c: https://spyglass.hhmi.2i2c.cloud/.

## Introduction

A central goal of neuroscience is to understand how the structure and dynamics of neural activity relate to the internal states of the organism and the external world. This understanding is derived from analyzing complex, multi-modal datasets. While the community has improved tools and algorithms for data collection and analysis^1–6^, extracting consistent and reproducible insights from data remains a complex and time-consuming task. Often, researchers take years to collect and organize data, which is then transformed through a complicated series of analyses using custom scripts. This begins with preprocessing that isolates specific signals from the data, followed by multiple subsequent analyses that quantify properties of these signals. The outputs of these analyses are then synthesized across datasets, and when they are consistent upon limited replication, they are reported in the scientific literature—with the data and analysis scripts documented to varying degrees.

Ideally, it would be possible for another group to take the same raw datasets, apply the analyses, and rapidly and reliably reproduce the findings. In practice, this is often exceptionally challenging. Raw data are seldom shared and metadata critical for understanding the data are often not included, posing a significant challenge to replication. Essential components of the analysis pipelines, such as the manual curation of sorted spikes and artifact rejection, are often irretrievable from the written reports. Similarly, the full set of parameters used for each of the analyses are not shared or hidden in cryptic analysis scripts. Efforts to reproduce findings are also hampered by idiosyncratic data and code organization, poor documentation, and missing vital details, including computational hardware requirements^7^. In collaborations among multiple scientists, these problems are exacerbated due to the variability in how each participant carries out analysis. Consequently, the full validation of a result usually requires repeating the experiment and reconstructing the analysis from scratch.

These difficulties in replication of analyses incur significant costs in time and effort. A new trainee might struggle to analyze existing data because they do not understand critical details. A scientist who downloads the data from a previous study may find that the analyses they wanted to carry out are impossible because the raw data is not available. Alternatively, raw data may be available, but the scientist may need intermediate results (e.g. spike waveforms) that are not included. Similarly, shared code, including visualizations, is most often not standardized or documented, causing multiple teams to duplicate efforts and implement the same tools.

A system that addresses these challenges therefore should enable:

- compilation of raw data with sufficient metadata for analysis and reuse
- sharing of data and all intermediate analysis results in an accessible format
- reproducible analysis via well-documented, organized, and searchable pipelines
- generation of shareable visualizations to facilitate communication and collaboration
- easy use by scientists with minimal formal training in data management.

Achieving these goals would represent a major step towards meeting the FAIR guiding principles for findable, <u>a</u>ccessible, interoperable, and reusable^8^ data and analysis pipelines^9^. For example, it would become possible to easily find publicly available data, analyze it with a standardized pipeline that keeps track of all the parameters, and generate a visualization to share the results over the web—a stark contrast to how science is practiced today.

In pursuit of this vision, many organizations, such as the Allen Institute for Brain Science (AIBS), Johns Hopkins Applied Physics Lab (APL), and the International Brain Laboratory (IBL), have made strides by standardizing and sharing data and analyses^10–12^. However, these efforts have not fully resolved the issues related to data sharing and reproducible analysis. For instance, a lack of raw data often precludes reproduction of early stages of the analysis, and many steps of the processing pipeline (e.g. the criteria used for manual or automatic curation of spike sorting) that can significantly affect the results^13^ are omitted. Some of the technologies used, such as cloud computing services (e.g., CodeOcean) and sophisticated databases and APIs^11,14,15^, can be cost prohibitive or require specialized software engineering expertise that is beyond the reach of most labs. Furthermore, these existing efforts tend to be focused on the needs of specific projects, data types, and behavioral paradigms, limiting their scope. Thus, while these efforts mark important advances, there remains a need for user-friendly, integrated solutions that can be widely adopted across individual labs in the neuroscience community.

To address this need, we developed Spyglass, an open-source neuroscience data management and analysis framework written in Python. Spyglass builds on widely available community-developed tools and adopts the Neurodata Without Borders (NWB) as the standardized format^16,17^. It uses DataJoint^5,18^ to manage reproducible analysis pipelines with a relational database and incorporates novel software tools (e.g. Kachery and Figurl) for sharing data and web-based visualizations to enable collaboration within and across labs. This includes methods for exporting and uploading all raw data and intermediate results used to produce a manuscript, which, along with sharing of code, enables full replication of results. Spyglass is Python-based and thus can accommodate pipelines that use a wide array of analysis packages that have been developed by the community, including SpikeInterface^19^, GhostiPy^20^, DeepLabCut^2^, and Pynapple^21^. Spyglass also offers ready-to-use pipelines for analyzing behavior and electrophysiological data, including spectral analysis of local field potential (LFP), spike sorting, video processing to extract position, and decoding neural data. Spyglass can be extended to support additional pipelines for behavioral, intracellular, optical physiology data, or other data types that can be stored in the NWB format. In addition to extensive documentation and tutorials, new users can try out a demo version of Spyglass hosted on the web by HHMI and 2i2c as a Jupyter Hub instance. Here we describe the structure of Spyglass and demonstrate its potential by applying the same analysis pipelines to NWB files from different labs and comparing the results.

## Results

### Overview of Spyglass

Spyglass is an open-source Python-based software framework for reproducible analysis of neuroscience data and sharing of the results with collaborators and the broader community (Figure 1). It is designed to be used by everyone in a laboratory who works with the data, both as a general-purpose tool to enable the development of new analysis pipelines and a tool that allows those pipelines and associated results to be frozen and packaged to enable reproducibility. It can be run locally or in the cloud. Analyzing data with Spyglass begins with raw data and experimental metadata stored in the NWB format^16,22^. These NWB files are ingested into a relational database and processed using DataJoint-enabled pipelines. Existing pipelines are built around common neuroscience tasks such as spectral filtering, spike sorting, pose tracking, and neural decoding and each user can extend these pipelines to carry out the specific sets of analyses needed for their project. DataJoint stores parameters of each analysis and tracks the intermediate analysis results, which are also stored as NWB files to maintain a shareable standardized data format. Spyglass enables sharing results and interactive visualization of the data over the web via Kachery and Figurl. Finally, Spyglass supports exporting specific parts of the database required to reproduce the results and figures of a manuscript and the upload of the associated raw data and analysis outputs to a public repository. In the following sections, we provide detailed descriptions of these components and the design decisions behind them.

**Figure 1:**
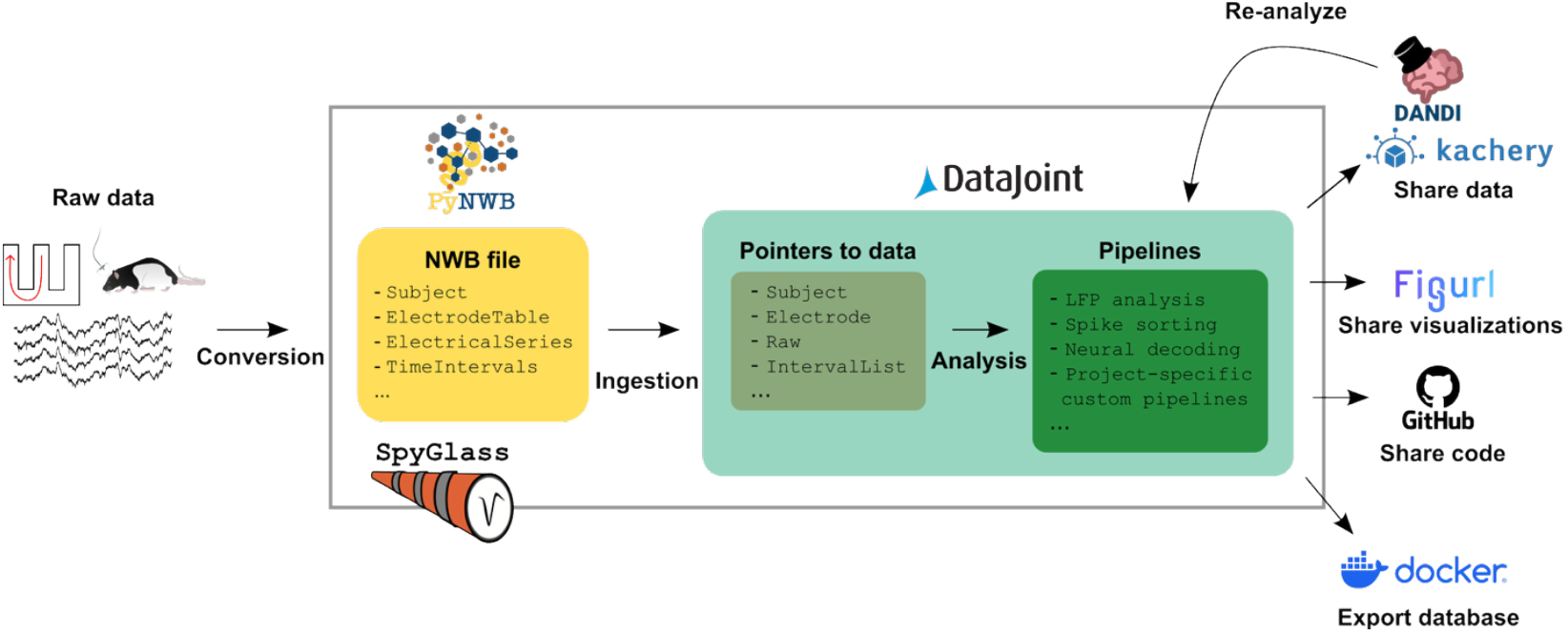
Overview of Spyglass. The raw data—consisting of information about the animal, the behavioral task, the neurophysiological data, etc.—is converted to the NWB format (yellow box) and ingested into the Spyglass database. The pipelines (dark green box) operate on pointers to specific data objects in the NWB file (tan box). The raw and processed data are then shared with the community by depositing them to public archives like DANDI or shared with collaborators via Kachery. Visualizations of key analysis steps can be shared over the web via Figurl. Code is shared by hosting the codebase for Spyglass and project-specific pipelines on online repositories like GitHub. Finally, the populated database may be shared by exporting it to a Docker container.

### The NWB format as data specification

#### Why NWB?

A typical neuroscience experiment consists of multiple data streams stored in different formats. Managing such heterogeneous data in a shareable and accessible manner is challenging. A practical solution is to save the data in a community-supported format like NWB, which is emerging as a standard for neurophysiology and behavior data^16,22^. We have chosen NWB as the data specification in Spyglass for the following reasons:

- The versatility of NWB accommodates various data types and allows metadata to be saved with the data in a single self-annotated file.
- NWB files are immediately shareable.
- Public data archives like DANDI^23–26^ accept the NWB format and provide [APIs to easily stream file contents](https://docs.dandiarchive.org/api/rest-api/) for local analysis.
- [Tools developed for NWB files](https://nwb-overview.readthedocs.io/en/latest/tools/analysis_tools_home.html) are immediately accessible to users.

Conversion to NWB can be done using software tools developed by the community, such as the [NeuroConv](https://github.com/catalystneuro/neuroconv) package or [NWB GUIDE](https://nwb-guide.readthedocs.io/), a desktop app for converting data to NWB without having to write code.

Importantly, Spyglass requires all raw data—including neurophysiology, behavioral task, interaction with the environment— to be in the NWB format *prior to any analysis*. This ensures reproducibility of all subsequent analyses by sharing the NWB file containing the raw data and the analysis pipelines. Furthermore, Spyglass stores virtually all intermediate results from downstream analysis pipelines in NWB. This ensures that all data associated with the analysis can be shared and read using the same software tools.

#### Spyglass-specific NWB requirements

NWB allows some flexibility in the specification of data to accommodate a broad range of experiments and lab-specific requirements. For example, the name of data types within the NWB file can differ from those expected by Spyglass. We have fully described the [Spyglass-specific NWB conventions](https://lorenfranklab.github.io/spyglass/latest/ForDevelopers/UsingNWB/) in the documentation website. To further accommodate NWB files from many sources, we have also developed a system that makes it possible to ingest NWB files into Spyglass even when they do not adhere to our naming conventions or best practices by including a configuration yaml file (see Methods and Table 1, 02_Insert_data).

**Table 1:**
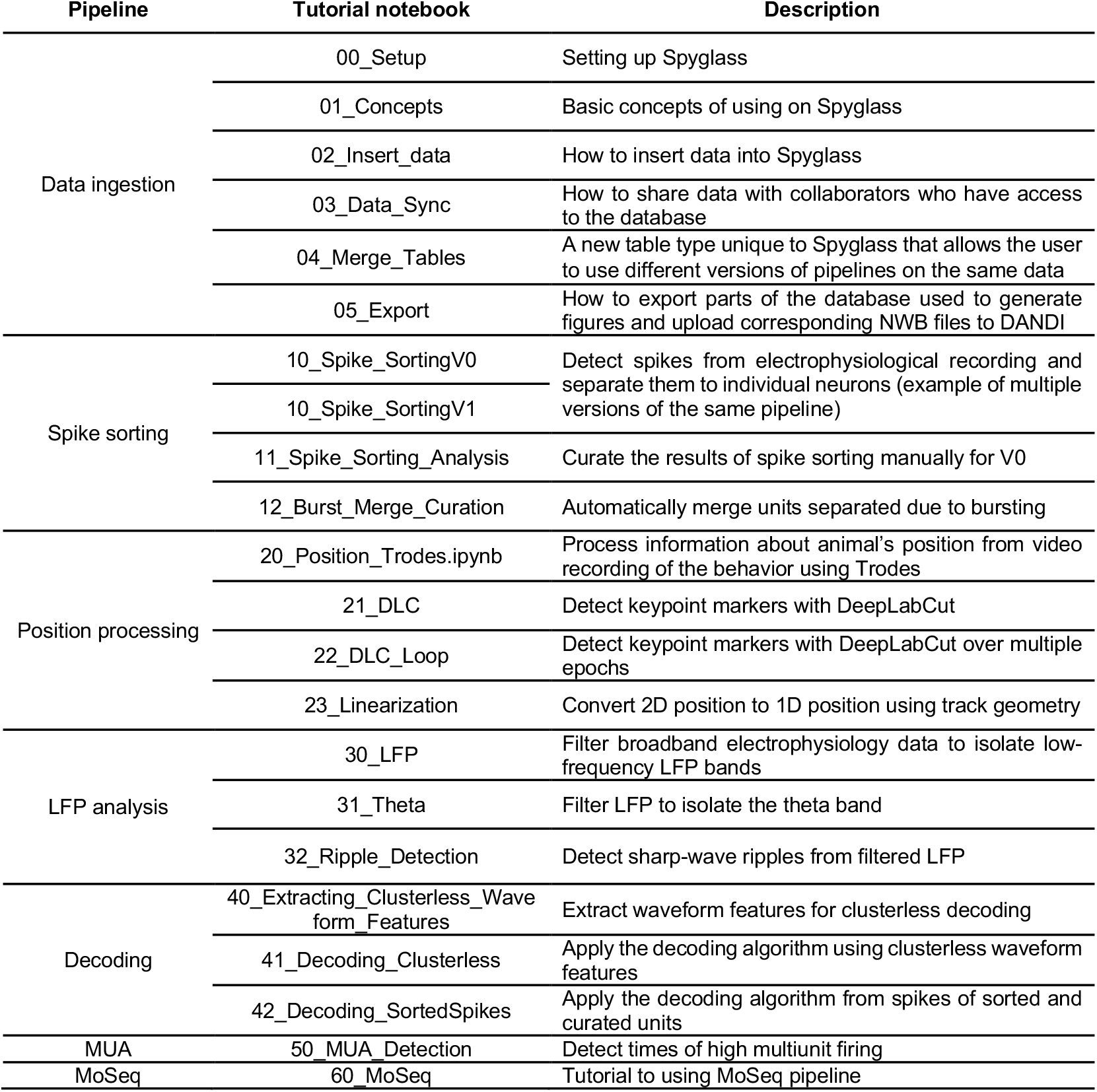
Tutorials included in Spyglass and their descriptions. All available from https://github.com/LorenFrankLab/spyglass.

### Relational database as analysis pipelines

#### Why a relational database?

One significant challenge with data analysis is in managing its complexity. Most results derive from an extended series of steps, including “preprocessing” (e.g. spike sorting for electrophysiological data, region-of-interest identification for optical physiological data, video processing for behavioral data, etc.) and downstream analyses. Each step depends on a different algorithm with a specific set of parameters and generates distinct intermediate data. Tracking these numerous components is difficult, and understanding how another scientist has managed them can be even more daunting. This complexity hinders collaboration, verification of results, and data reuse.

These issues motivated our use of a formal software system: the relational database, a well-established data structure that uses tables to organize data. To construct an analysis pipeline, we use DataJoint^27,28^ to define a series of database tables with a dependency structure. Associated with the database tables is code that carries out an analysis using a specific set of parameters and a specific part of the data. This code then stores the results as a new row in the table with a pointer to the results stored on disk as an NWB file. Creating a new table row is referred to as “populating” the table. Thus, data analysis becomes a matter of populating and interacting with the database. This style of data analysis offers many advantages:

- It lowers effort for users seeking to apply the same analysis to multiple datasets, as they only need to specify the data and parameters for computation (“what”) independent of the execution details (“how”).
- It provides a structure to organize and systematize the analysis parameters, data, and outputs into different tables. This contrasts with user-generated configuration files where each user could adopt their own idiosyncratic approach to specifying parameters and data.
- It enables easy access to multiple datasets via queries (e.g. to find all datasets with recordings from a particular brain region or that used a particular behavioral paradigm).
- It is concurrently accessible to multiple users.

Because DataJoint binds the code for running the computation with the table that will store the result, populating the same table will execute the same code. DataJoint also provides additional features for reproducible data analysis, such as maintaining data integrity of the database (e.g. deleting a table entry causes cascading deletion of dependent entries in downstream tables) and the files containing the results (e.g. by checksum verification).

#### How does Spyglass differ from DataJoint?

While Spyglass is based on DataJoint, it offers many useful features that DataJoint lacks. These include:

- A tight integration with the NWB format: When the NWB file is ingested into Spyglass, pointers to the data types appear as rows in a set of predefined tables. These serve as the starting point for analyses and an interface for the users to access the raw data within Spyglass. We provide the [mapping between NWB data types and corresponding Spyglass tables](https://lorenfranklab.github.io/spyglass/latest/ForDevelopers/UsingNWB/#object-table-mappings) in the documentation website.
- Extending table types: Spyglass provides a mix-in class, which allows different tables to inherit shared behaviors without duplicating code, for defining table types that are not included in DataJoint or extending the function of existing table types. This is used to implement many key table types such as `Merge` tables, which allow multiple upstream pipelines to feed into the same downstream pipeline. This example is illustrated in the description of the spike sorting pipeline below.
- Permission-based delete: Spyglass enables the deletion of individual rows in a table based on pre-defined user permission. This is not naturally supported by MySQL, the underlying relational database management system used by DataJoint.
- Improved searching based on restrictions on non-primary keys: Spyglass allows the users to conveniently track the provenance of a particular row in a downstream table across multiple upstream tables with only partial information.
- Export system for publishing: Spyglass provides a convenient way to export only the part of the database used for generating results and figures for a publication. This is done by caching the information about tables that are accessed when generating figures.
- The inclusion of various helper functions, which are detailed on the Spyglass documentation website.

### Setup and hardware requirement

Spyglass can be installed in any setting that can support Python via the Python Package Index (PyPI). We provide detailed installation instructions on the documentation website, including a complete list of software requirements. In addition to the Python package, using Spyglass requires running a relational database (currently MySQL backend is supported). In our laboratory, we run this from a Docker container provided by DataJoint on a lab-wide server and grant access to it to members of the lab and other collaborators. This local configuration is recommended for use cases involving ∼ 10 users. For a larger scale deployment, one could also run the Spyglass database in the cloud using services such as AWS.

### Practical use cases and extensions

Spyglass comes with many pre-defined pipelines that implement common analysis tasks for electrophysiological and behavioral data. For users interested in using these pipelines, they can do so as soon as they ingest their NWB files into the database. Spyglass can also serve as a jumping off point for exploratory data analysis. For example, the user can conveniently read specific data types from the NWB file by first ingesting it into Spyglass and accessing database tables with Spyglass functions (e.g. `fetch_nwb`) or load those objects in a format compatible with Pynapple^21^ (`fetch_pynapple`). If they need to pre-process the data first, they can do so by running the relevant pipelines. Once the user has decided to formalize a particular analysis that is not yet supported by Spyglass, they can extend Spyglass and create user-generated custom pipelines. These could include data types from NWB files not currently supported by Spyglass (e.g. photometry, optical physiology, etc.) or build on existing Spyglass pipelines. Because the raw data and intermediate results are in NWB format, the custom pipelines can take advantage of analysis software packages within the NWB ecosystem.

### Organization of analysis pipelines

Here we delve deeper into the design and organization of analysis pipelines in Spyglass. As mentioned previously, the analysis pipelines are defined as a set of tables in the relational database. Specifically, Spyglass uses DataJoint syntax to define tables as Python classes (see online documentation on [Custom Pipelines](https://github.com/LorenFrankLab/spyglass/blob/master/docs/src/ForDevelopers/CustomPipelines.md) and this [video](https://youtu.be/-2R7HLmmyqs) for examples). The code for executing the analysis is associated with these tables as class methods, enabling a tight integration of the database structure with the code for populating it. We refer the reader to the DataJoint documentation for more details on specific commands to interact with the database.

When an NWB file is first ingested into Spyglass, pointers to the data types in the NWB file are stored in database tables of the `Common` module. Each `Common` table corresponds to a data object in the NWB file and serves as an interface to retrieve it with simple function calls (`fetch_nwb`). The retrieval is “lazy” in the sense that only a specific part of the data is loaded for analysis instead of the entire NWB file.

An analysis pipeline consists of sets of tables downstream of the `Common` tables. In each step in the analysis, the user populates one of four table types (Figure 2A):

**Figure 2:**
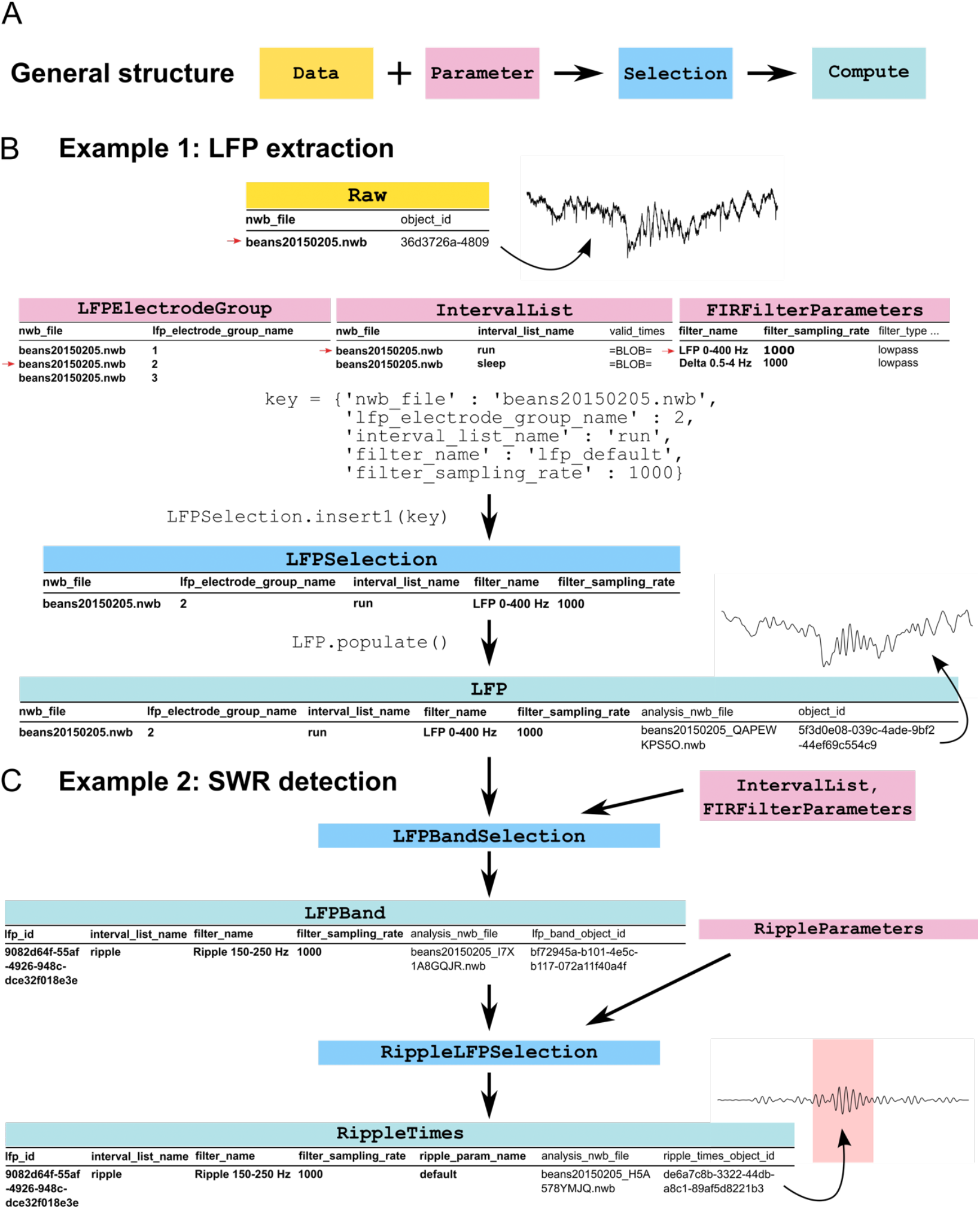
Analysis pipelines in Spyglass. (A) A general structure for a Spyglass pipeline. (B) Example 1: LFP extraction. Note the correspondence to the pipeline structure in (A) as shown by the color scheme. The trace next to the `Raw` table is raw voltage data sampled at 30 kHz and is represented by a row in the `Raw` table. This, along with parameters from `LFPElectrodeGroup`, `IntervalList`, and `FIRFilterParameters` tables (red arrow), are defined in a Python dictionary and the `LFPSelection.insert()` call is used to insert the reference to the raw data and the now associated parameters into `LFPSelection` table. When the `populate` method is called on the `LFP` table, the filtering is initiated and the output is inserted into the database. The results (e.g. the trace above `LFP` table) are stored in NWB format and its object ID within the file is also stored as a row in `LFP` table, enabling easy retrieval. (C) Example 2: Sharp-wave ripple (SWR) detection. Note that the key specification, insert, and populate calls are omitted for simplicity. This pipeline is downstream of the LFP extraction pipeline and consists of two steps: (i) further extraction of a frequency band for SWR (`LFPBand`); and (ii) detection of SWR events in that band (`RippleTimes`). Note that the output of LFP extraction serves as the input data for the SWR detection pipeline and can thus be thought of as both `Compute` and `Data` types. As in (B), for each step, the results are saved in NWB files and the object ID of the analysis result within the NWB file are stored as rows in the corresponding `Compute` tables. The trace above the `RippleTimes` table is the SWR-filtered LFP around the time of a single SWR event (pink shade). In each table, columns in bold are the primary keys. Arrows depict dependency structure within the pipeline.

1. `Data` tables contain pointers to data objects in either the original NWB file or ones generated by an upstream analysis.
2. `Parameter` tables contain a list of the parameters needed to fully specify the desired analysis.
3. `Selection` tables allow users to select and pair a data entry and a parameter entry, defining the input to the `Compute` table.
4. `Compute` tables execute the computations to carry out the analysis using the `Data` and `Parameters` specified in the `Selection` table entry. These results are then stored and can serve as `Data` for downstream analysis.

This design has multiple features that we have found to be beneficial. First, `Parameter` tables store the full set of parameters needed to specify a given analysis. For example, a `Parameter` table entry for a firing rate analysis of a single neuron might specify the bin size and smoothing to be used for that analysis. Multiple such entries can be defined, allowing a user to select the most appropriate one for the question being addressed. Second, because `Selection` tables specify which `Parameter` table entry was used for a given analysis on the associated `Data` table entry, they provide the key information needed to know which parameters were used to generate the entry in the downstream `Compute` table. Third, it is simple to associate a given `Data` table entry with multiple `Parameter` table entries and then re-run the analysis on those pairs. This enables a user to understand how their choice of parameters impacts their results, something that is otherwise difficult to manage and track.

Spyglass includes pipelines for a diverse range of analysis tasks in systems neuroscience, such as the analysis of LFP, spike sorting, video and position processing, and fitting state-space models for decoding neural data. Tutorials for all pipelines are available on the [Spyglass documentation website](https://lorenfranklab.github.io/spyglass/latest/) (Table 1). Our goal was take advantage of other open source packages, and we have therefore integrated support for Pynapple^21^, a general purpose neural data analysis package. We also built our pipelines to take advantage of other community-developed, open-source packages, like GhostiPy^20^, SpikeInterface^19^, DeepLabCut^2^ and Moseq^29^. These pipelines store a complete record of the analysis and simplify the application of these tools. Furthermore, multiple versions of the pipelines can co-exist to apply different algorithms to a single data set, making it easy to probe the robustness of the results (see *Merge motif* below). Finally, the pipelines are modular as long as they process different kinds of data stored in the NWB files.

Next we provide a detailed description about the implementation of three common analysis tasks in Spyglass pipelines: (i) filtering broadband extracellular voltage traces to extract the lower-frequency LFP bands; (ii) detecting discrete events (e.g. sharp-wave ripples, a hippocampal event marking the time of bursts of population activity) in the LFP signals; and (iii) spike sorting and curation.

#### Example 1: LFP extraction (Figure 2B)

To extract the LFP signal (below 400 Hz), we use the pipeline shown in Figure 2B. First, we select a row from the `Raw` table, a `Data` table that points to an `ElectricalSeries` object in the NWB file. We then specify the parameters of the analysis in the `Parameter` tables: the list of channels for which LFP should be extracted (`LFPElectrodeGroup`), the time interval for the LFP extraction (`IntervalList`), and the coefficients for the filter that will be used on the data (`FIRFilterParameters`). These parameters are associated with the entry in the `Raw` table by defining a Python dictionary object that specifies the `Data` and `Parameter` entries and inserting it into a `Selection` table (`LFPSelection`) by calling the `LFPSelection.insert1` method (Figure 2B). Finally, we apply the filter to the selected data over the selected interval using the `LFP` table (a `Compute` table) by calling the `LFP.populate` method. The resulting filtered data is saved to disk in the NWB format, and the object ID associated with the LFP object within the NWB file is also stored in the `LFP` table for easy retrieval. Thus, the corresponding entry in the `LFP` table contains all the details about the data and the parameters, allowing a user to fully track the provenance of the output.

#### Example 2: Sharp-wave ripple detection (Figure 2C)

Once the LFP extraction is completed, we can build on the results by applying another filter to isolate a specific frequency band and identifying sharp-wave ripples (SWRs), a prominent LFP event within hippocampal data. This pipeline is illustrated in Figure 2C. It applies two additional steps to a row in the `LFP` table: another band-pass filter to isolate the 150-250 Hz band and a subsequent detection of SWR events. Each step uses the same basic scheme shown in Figure 2A. These include defining a specific band-pass filter in the `Parameter` tables; selecting a time interval for the bandpass filtering; and adding an entry to `LFPBandSelection` table that binds both the filter parameters and the time interval with a row in the `LFP` table. A call to `LFPBand.populate` generates an NWB file containing the ripple-band data and an entry in the `LFPBand` table with information about which data and parameters were used. Next, the user selects an entry in `RippleParameters` to define the parameters for detecting the ripple events (e.g. threshold over the spectral power) and associates it with filtered data in `LFPBand` in the `RippleLFPSelection` table. Finally, the `RippleTimes` table is populated (by `RippleTimes.populate)`, which identifies the start and end times of each ripple event and saves these to a new NWB file.

#### Example 3: Spike sorting and curation (Figure 3)

The spike sorting pipeline (Figure 3) combines the principles of analysis pipeline design we outlined previously with additional design features. This pipeline uses SpikeInterface^19^ to perform the operations critical for spike sorting, but also tracks all of the parameters used and provides a system for tracking multiple sorting curations. The pipeline includes the following steps: (1) preprocess the recording (e.g. filter and whiten to remove noise); (2) apply spike sorting algorithm (e.g. MountainSort4, Kilosort3, etc.); (3) curate the results (e.g. either manually or automatically by computing quality metrics); and (4) consolidate the output with other sources of sorted units (e.g. those already present in the NWB file) for downstream analysis. Each of these steps follow the general design shown in Figure 2A. We also detail additional features that have not been discussed previously.

**Figure 3:**
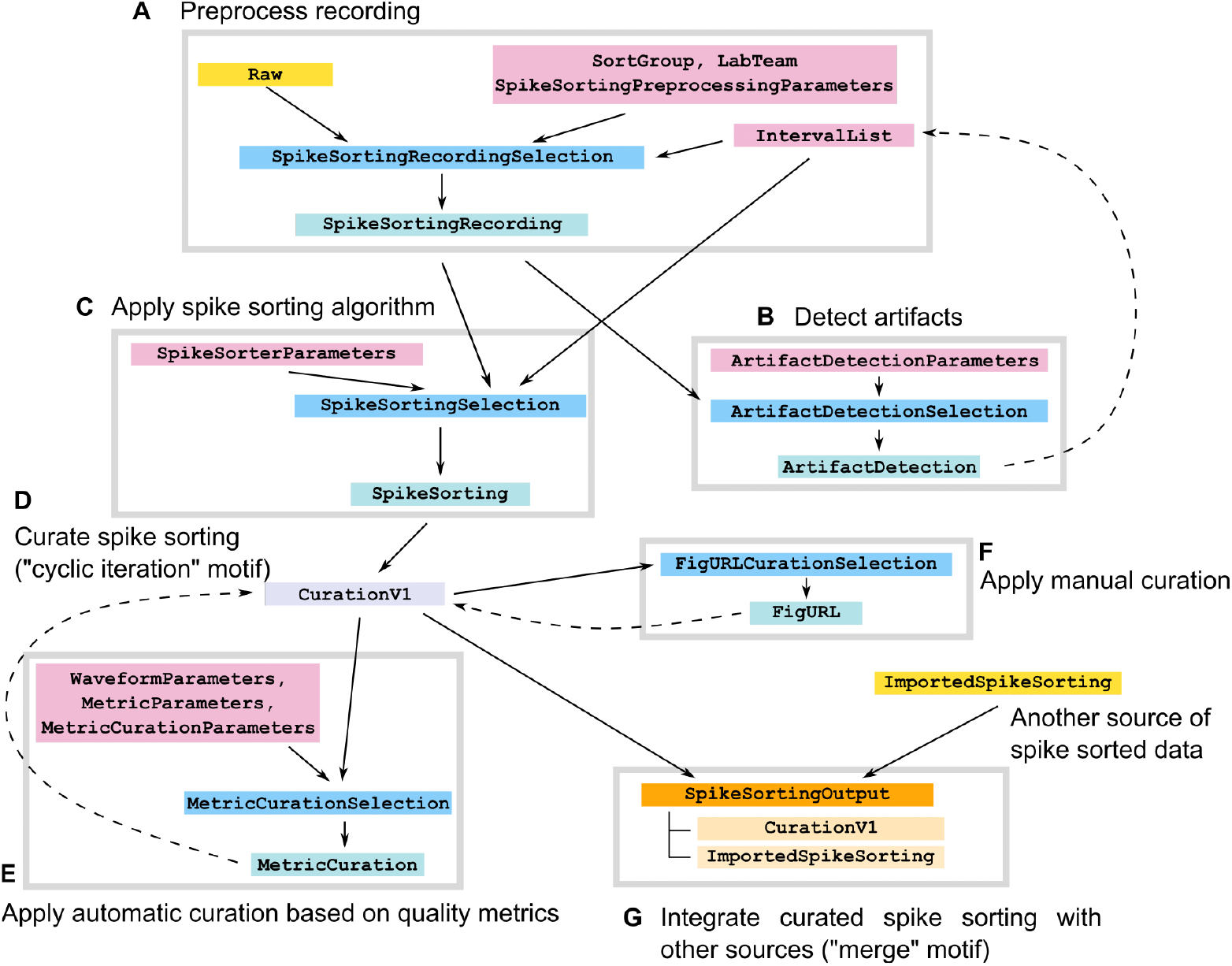
Spike sorting pipeline. The Spyglass spike sorting pipeline consists of seven components (large gray boxes), all of which take advantage of the SpikeInterface library: preprocess recording (A); detect artifacts to omit from sorting (B); apply spike sorting algorithm (C); curate spike sorting (D), either with quality metrics (E) or manually (F); and merge with other sources of spike sorting for downstream processing (G). Solid arrows describe dependency relationships and dashed arrows indicate that the data is re-inserted upstream for iterative processing. Note the two design motifs (see text): “cyclic iteration” for curation and “merge” for consolidating data streams. Color scheme is the same as Figure 2, except for light purple (cyclic iteration table), orange (merge table), and peach (`Parts` table of the merge table).

##### Global Parameter tables (e.g. IntervalList)

An important object in any analysis is the time interval during which the data were collected or to which analysis procedures should be applied. To avoid having a separate table for time intervals in every pipeline, we store them in the `IntervalList` table of the `Common` module for all pipelines. For example, in the spike sorting pipeline (Figure 3), `IntervalList` provides a time interval for both preprocessing the recording (`SpikeSortingRecordingSelection`) and running a spike sorting algorithm (`SpikeSortingSelection`). In addition, the intervals during which artifacts (i.e. high-amplitude voltage transients from behavioral events such as licking) occur is identified and fed back into `IntervalList` (dashed arrow in Figure 3).

##### “Cyclic iteration” motif for curation

Certain pipelines, such as curating the output of spike sorting, may need to be run multiple times on the same data. For example, one might first compute quality metrics to identify noise clusters and potential candidates for merging over-clustered units (Automatic); then inspect, merge, and apply curation labels to the result with an external viewer (Manual); and finally, compute a final set of metrics to describe the quality of each unit (Automatic). This results in a sequence of curation steps: Automatic, Manual, Automatic. Depending on the data, the user may choose a different curation sequence, and the order and length of these sequences might change as new algorithms and metrics are developed. This presents a challenge in modeling the pipeline within the relational database.

We therefore developed a specific design motif to enable this iterative curation with a finite number of tables (Figure 3). First, a given row of the `CurationV1` table (the output of the spike sorting step) is taken through automatic or manual curation steps downstream. Upon completion, the spike sorting object may enter this curation pipeline again as a new row in the `CurationV1` table. Importantly, the new row has information about previous curation from which it descended. This allows the user to track each round of curation while applying as many steps as desired. It can also be easily extended; if new automatic curation algorithms are developed in the future, it can simply be added downstream to the `CurationV1` table, enabling application of the latest methods to previously collected data.

##### “Merge” motif for consolidating data streams and versioning pipelines

A different challenge arises when the user wants to feed multiple streams of data of the same type into a single downstream pipeline. For example, once curation is completed, the spike sorting is saved in `CurationV1`. But some NWB files may already contain curated spike sorting (as a row in the table `ImportedSpikeSorting`), and one may want to apply the same downstream pipeline to both data sources to compare the results. In yet another case, the other data stream could be a different version of the spike sorting pipeline (e.g. `CurationV2`) that uses different algorithms but produces output of the same type. Adding the same downstream pipeline to each of these separately would result in code redundancy and database bloat. Simply having these converge onto a single downstream table is not desirable either, as it will require modifying an existing table to add new columns every time a new version or new data stream is added.

To solve this problem, we have designed a “merge” table type (Figure 3). Here `Parts` tables (a table type within DataJoint tightly associated with a parent table) are used to implement the merging of multiple data streams onto a single table. The downstream pipeline then gets data from this table without any duplication. More details for the implementation and helper functions to maintain data integrity can be found in the tutorial notebook (Table 1, 04_Merge_Tables).

### Sharing Data, Analysis, and Visualization

#### Complete sharing of data and analysis at the end of projects

A key goal of our system is to simplify sharing data and analyses when results are ready to be published. Because all raw and intermediate data are in the NWB format, they can be directly deposited to DANDI^16,24–26^, a NIH-supported public archive for neuroscience data. Sharing the analysis code is also easy: simply share the codebase for the analysis pipelines (i.e. Spyglass plus any project-specific pipelines) and the scripts used to populate the database. Others can then download the raw data from DANDI, set up the database with Spyglass, and recreate all results locally by executing the population script. Alternatively, users may want to share the Spyglass database in its populated state so that the community can access it directly without going through the setup procedures or re-running time-consuming analysis steps. This can be done by (i) hosting the database on the cloud and granting access to users outside the lab; or (ii) exporting and sharing parts of the database that were used by the project. Spyglass facilitates the second option by providing functions that automatically log the table entries and NWB files used for creating figures of a manuscript in a Python environment (Table 1, 05_Export). The dependencies of these entries are traced through the database to compile the complete set of raw, intermediate, and plotted NWB files and their corresponding database entries. These are stored in the `Export` table, which also generates a bash script to create SQL dumps of the identified database entries.

To upload these files to DANDI, users must first [register a new dandiset](https://docs.dandiarchive.org/user-guide-sharing/creating-dandiset/) for their project and record their API and dandiset ID. With this information, they can then use the method `DandiPath.compile_dandiset()` to automatically validate, organize, and upload all project files to the DANDI archive. Additionally, this process stores the archive information for each file in the `DandiPath` table, allowing `fetch_nwb` to automatically stream data from the DANDI cloud storage when not available locally.

To create a sharable Docker image of the project, we provide a template repository called spyglass-export-docker. Users first download a local copy of this repo and copy the SQL dump file, environment yaml, and figure-generating notebooks generated during Spyglass export into the appropriate folders. Running the provided `docker-compose` scripts then generates two linked Docker containers: one running the reconstructed Spyglass SQL database, and a second connected to this database and running a Jupyter Hub—with a python environment matching that used when generating the figures. These can be readily shared with new users to provide them immediate access to all steps of the analysis process and the corresponding data through DANDI streaming

#### Controlled sharing for ongoing projects

For ongoing projects, users may want to limit the sharing of the analyses to their collaborators. This requires controlling access to the database and the underlying NWB files that contain the raw or intermediate data. This is straightforward to manage in Spyglass. DataJoint handles access to the database natively by requiring a username and a password. Managing access to the NWB files is handled by [Kachery](https://github.com/flatironinstitute/kachery-cloud), a content-addressed sharing tool for scientific data (Figure 4A). Specifically, the user selects the NWB files to be shared by inserting pointers to them into `NwbKachery` and `AnalysisNwbKachery` tables within Spyglass. When the collaborator attempts to access these files, Kachery first looks for them in their local system. If not found, the corresponding files are automatically uploaded from the user’s system to a cloud storage server and then downloaded to the collaborator’s computer. This feature is detailed in a tutorial (Table 1, 03_Data_Sync). Critically, the downloaded files are never modified locally within Spyglass, and attempt to access a modified file would result in a DataJoint error. This ensures that each user is working on the same underlying data even if they are at different sites. More generally, Kachery offers advantages over other file hosting services (e.g. Dropbox and Google Drive) or alternative architectures (e.g. IBL data architecture) by not requiring a central location to track available files and providing a user-friendly Python API. We point interested readers to the [Kachery GitHub repo](https://github.com/magland/kachery) for further descriptions.

**Figure 4:**
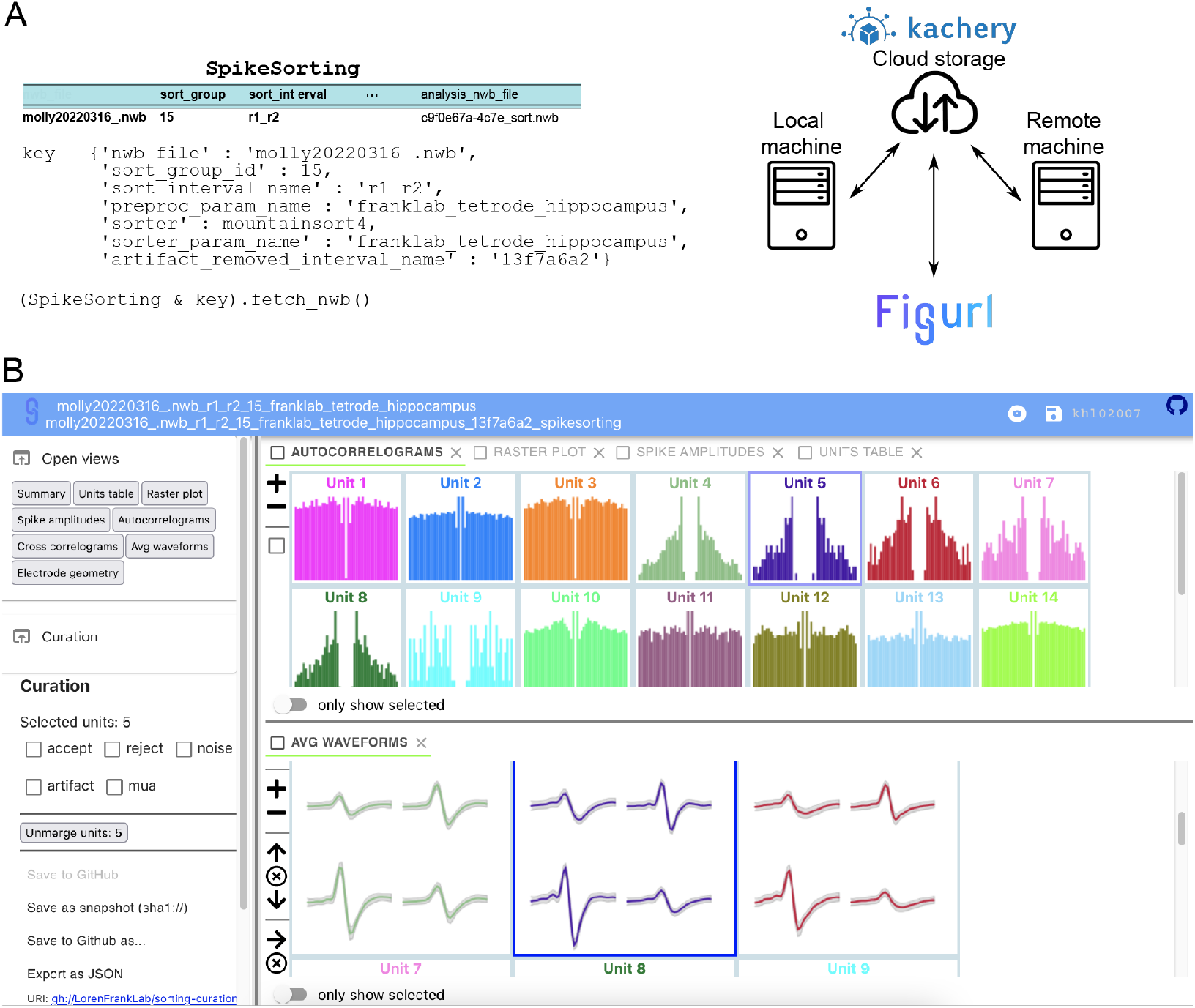
Sharing data and visualizations. (A) Kachery provides a convenient Python API to share data over a content-addressable cloud storage network. To retrieve data from a collaborator’s Spyglass database, one can make a simple function call (`fetch_nwb`) that pulls the data from a node in the Kachery Zone to the local machine. (B) Example of a Figurl interactive figure for visualizing and applying curation labels to spike sorting over the web.

#### Sharing visualizations

Spyglass enables users to create and share interactive visualizations of final and intermediate analysis results through the [Figurl](https://github.com/flatironinstitute/figurl) package. These visualizations facilitate understanding complex, multi-modal neuroscience datasets by allowing users to (i) quickly compare different stages of processed data to spot issues with their data and (ii) align multimodal information sources to get a more holistic view of their dataset. Figurl is integrated within Spyglass as dedicated tables attached to specific pipelines such as spike sorting (Figure 4B) and neural decoding (Figure 5). Populating these tables generates a URL to web-based visualizations for exploring complex, multi-dimensional time series across multiple views whose time axes can be linked. Sharing them is also easy, as the URL can be accessed from any browser without the need for local software installation or specialized hardware. This allows collaborators anywhere in the world to easily access and explore the data.

**Figure 5:**
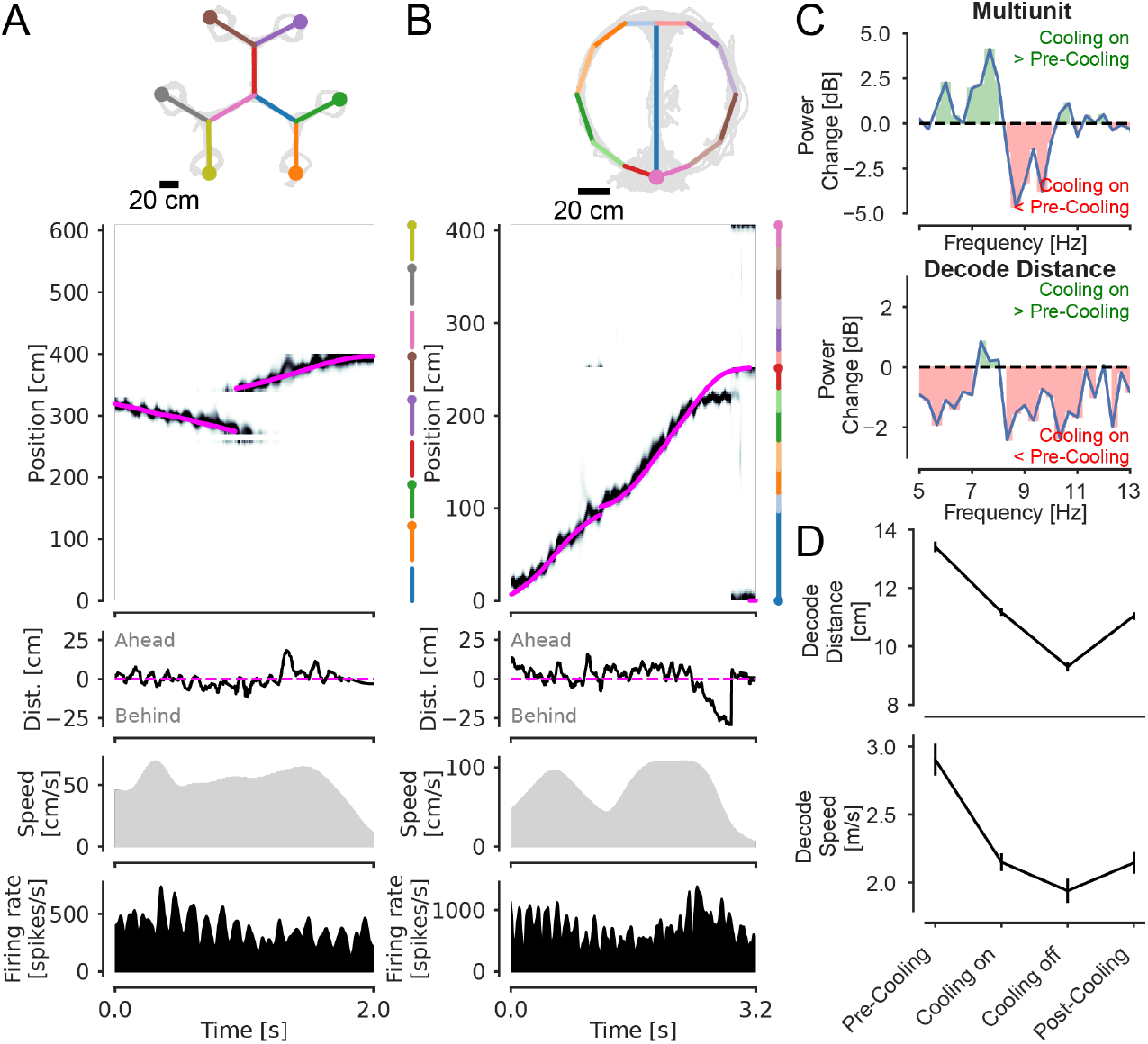
Applying decoding pipelines to multiple data sets from different labs. (A) Decoding neural position from rat hippocampal CA1 using a clusterless state space model (UCSF dataset). In the top panel, grey lines represent positions the rat has occupied in the spatial environment. Overlayed lines in color are the track segments used to linearize position for decoding. Filled circles represent reward wells. The second panel from the top shows the posterior probability of the latent neural position over time. The magenta line represents the animal’s actual position. The vertical lines on the right represent the linearized track segments with the colors corresponding to the top panel. The third panel from the top shows the distance of the most likely decoded position from the animal’s actual position and sign indicates the direction relative to the animal’s head position. The fourth panel from the top is the animal’s speed. The final panel is the multiunit firing rate. (B) Decoding from rat hippocampal CA1 using existing spike sorted units (NYU dataset). Conventions are the same as in A. Filled circle in the linearization represents the reward zone rather than the reward well. (C) Decoding analysis of the NYU dataset. The top panel shows the power difference of the multiunit firing rate between the medial septal cooling period and the pre-cooling period in the 5-13 Hz range. The power at 8-10 Hz is attenuated during cooling while the power at 5-8 Hz is enhanced, showing a slowing of the theta rhythm during cooling. The bottom panel shows that the power of the distance between decoded and actual position (decode distance) is mostly reduced throughout the 5-13 Hz range. (D) Cooling decreases the decode distance and speed and this effect may only recover partially after cooling. Bars represent 95% confidence intervals.

### Demonstration of generalizability: neural decoding of position in multiple data sets

A major goal of Spyglass is to facilitate the analysis of data across multiple datasets that may come from different laboratories. To illustrate this, we ingested and analyzed two NWB files containing single-neuron recordings from rat hippocampus, one from our laboratory and another from the Buzsáki laboratory at NYU^30^. Specifically, we applied a switching state space model ^31,32^ to decode the animal’s position from spikes and infer periods of different types of non-local representations (such as replay and theta sequences), during which the decoded position deviates from the animal’s true position. This is a complex analysis that involves integrating multiple data sources, including position and neural spiking activity, and applying an advanced statistical model with many user-defined parameters. The decoding pipeline in Spyglass enables the user to carry out every step of this analysis, including “preprocessing” of the data (e.g. linearize the 2D position of the animal, perform spike sorting, or import units that have already been sorted) and fitting of the model (see Supplementary Figure 1 for a visualization of the steps involved). After running the decoding pipeline, we visualize the results on the browser via Figurl and generate plots to further reproduce the reported results.

The UCSF dataset contains large-scale hippocampal recordings in a rat performing a foraging task in a maze with six reward sites and dynamic reward probabilities (Figure 5A, top panel). Applying the decoding pipeline to these data yields a probability distribution over space in 2 ms bins that describes our estimate of the “mental” position of the animal. This mental position tracks the animal as it traverses the maze (Figure 5A, 2^nd^ panel from top; see interactive visualization via Figurl) but also shows interesting systematic deviations from actual position. Computing the distance between the peak of the probability distribution and the actual location reveals characteristic patterns of such deviations from the actual position (Figure 5A, 3^rd^ panel from top) in which the decoded position sweeps ahead of the actual position and then back during movement bouts. This pattern recurs at ∼8 Hz, reflecting the well-known “theta sequences” seen in the hippocampus^33,34^.

We then applied this same pipeline to the NYU dataset, where rats performed a spatial alternation task on a maze with a figure-8 topology (Figure 5B, top panel). As expected, we could identify theta sequences in these data as well, highlighting the robustness of these phenomena (Figure 5B, 2^nd^ and 3^rd^ panels from top, see interactive visualization via Figurl). Moreover, the NYU dataset includes a specific manipulation in which the medial septum, a brain region critical for pacing the theta rhythm, was cooled, reducing the theta frequency from 8-10 Hz to 5-8 Hz. The authors originally carried out several detailed analyses to demonstrate that cooling reduced theta frequency and impaired behavior without changing the overall spatial tuning of single neurons or their tendency to fire sequentially within theta cycles. However, the authors did not apply state-space decoding methods, and did not characterize the effects of cooling on the decoded representation of space in relation to the animal’s actual position. We therefore applied our decoding pipeline to the cooling trials (“cooling on”) and the control trials preceding it (“pre-cooling”), just after it (“cooling off”), and the recovery trials 10-12 minutes after cooling (“post-cooling”).

The results of these analyses were consistent with the published findings and provided new characterizations that could serve as the foundation for additional discoveries. We first estimated the multiunit firing rate as a proxy for the theta LFP and characterized its power spectrum before and after cooling. As expected, cooling decreased the power above ∼8 Hz and increased the power below ∼8 Hz, consistent with the slowing of theta LFP shown in the original manuscript (Figure 5C, top panel). We then applied the same analysis described above to the distance between the decoded and the actual position during movement (“decode distance”), expecting cooling to have a similar effect on its power spectrum. Interestingly, here cooling led to a decrease in power at essentially all frequencies (Figure 5C, bottom panel). Consistent with this result, the decode distance decreased from the pre-cooling to cooling period, with a partial recovery during the post-cooling period (Figure 5D, top panel). Similarly, the average speed at which the decoded position moved ahead and behind the animal was also reduced during cooling and showed a partial recovery after the cooling period (Figure 5D, bottom panel). These results indicate that cooling reduces both the extent and the rate at which the decoded position deviates from the actual position. This was unexpected given that cooling had no effect on the average spatial tuning of these cells^30^. It also raises an interesting hypothesis: hippocampal representations of distant locations may be exquisitely tuned to the specific frequency of the rhythmic input from medial septum, such that slowing the rhythm down by just 2-3 Hz significantly limits their expression. More broadly, these findings illustrate the power of our framework that enables both replication of results across datasets and the re-analysis of previously collected data.

## Discussion

### Summary of results

Science is a social enterprise that relies heavily on collaboration and transparency among researchers^35,36^. Reproducible and shareable data analysis plays a critical role in this context, as it ensures that scientific findings can be independently verified and built upon by others. To facilitate this for the neuroscience community, we built Spyglass, a software framework that combines the NWB format and the relational database structure. Building on many community-developed tools, it provides useful features to design complex analysis pipelines, share raw and processed data, generate web-based visualizations, and analyze data from multiple sources. As a result, it simplifies collaboration within and across labs, making it well-suited as a community framework for neurophysiological and behavioral data analysis.

### Comparison to prior work

Our work builds on many previous approaches that have been proposed for scientific data management and reproducible analysis pipelines. This includes work from individual laboratories that have illustrated how a few elements of an NWB file could be read into a DataJoint database^37^, and publications highlighting datasets available in NWB^38^. More broadly, DataJoint is used by many labs with lab-specific pipelines^39^, but to our knowledge none of these efforts integrate cross-laboratory data and visualization tools or use NWB as the foundation to facilitate sharing. Our system also contains elements similar to those developed by large collaborative groups like The International Brain Laboratory (IBL) that are designed to organize neurophysiology data for sharing with collaborators and a module to automatically run analyses^12^. But the conversion to a standardized format (outside the collaboration or group) and public data sharing are only done following substantial analysis in the IBL system, complicating replication of the full analysis.

Other approaches do away with the relational database altogether. For example, DataLad uses version control tools such as git and git-annex to manage both code and data as files^40^. This enables the creation of a data analysis environment and decentralized data sharing. For building analysis pipelines, it may be combined with other tools for managing the sequential execution of scripts. For example, Snakemake^41^ (and related projects such as Cobrawap^42^) allows the users to gather and define the input, output, and the associated scripts to execute for each analysis step, thereby tracking the dependency between steps. But because these tools do not provide any formal structure for data analysis or parameter specification, they lack the advantages of the relational database that we discussed, such as being able to easily organize or search for the records of previous analysis based on specific parameters, efficient data sharing and access management to multiple users, and built-in data integrity checks based on constraints native to the database (e.g. primary keys).

By contrast, Spyglass begins with a shared data format that includes the raw data and offers both transparent data management and reproducible analysis pipelines using a formal data structure. One distinguishing feature of Spyglass is the emphasis placed on combinatorial matching of data and method in a reproducible way. For example, Spyglass makes it easy to apply multiple spike sorting algorithms to a given dataset and to compare the results, as this involves simply matching the data with different rows in the parameter tables. Spyglass also makes it straightforward to apply complex analyses like decoding to datasets from multiple labs, facilitating replication and data re-use. The system can be extended by adding new pipelines to the existing database as better tools and algorithms become available. These features enable the re-analysis of data to examine how the results depend on the choice of algorithm. We believe it is critical to provide this kind of future-compatibility to maximize the impact of the years of experimental work that go into each dataset.

### Limitations

Although Spyglass provides many useful features for reproducible data analysis, it has several limitations. Because of the central role played by the NWB format in Spyglass, a potential user must first convert their data to NWB, which requires time and effort^43^. In addition, some data types are yet to have defined standards within NWB (e.g. surgical procedure details, descriptions of conditions, detailed subject information), and if the user wishes to include those details, they would need to build an NWB extension and parallel Spyglass tables to do so. NWB also allows users to choose their own names for some datatypes (e.g. behavioral tasks), further requiring standardizations to agree on naming conventions.

In addition, users are expected to set up and maintain a relational database, which may involve additional training. Using Spyglass includes learning to work with the structure of DataJoint, such as the strict data integrity requirement that can make modification of existing tables difficult. Spyglass also does not yet include pipelines for processing certain types of neural data, such as optical physiology or fiber photometry, and some of its features such as Kachery-based file sharing may not currently support Windows (although it may be possible to run on the Windows Subsystem for Linux). Finally, as for all software frameworks, the evolution or lack of maintenance of other packages presents a challenge for long term support and reproducibility.

Fortunately, there are ongoing efforts to address these challenges. These include tools to simplify the raw data conversion into NWB, such as [NeuroConv](https://neuroconv.readthedocs.io/), a package to convert neurophysiology data in common formats to NWB automatically, and [NWB GUIDE](https://nwb-guide.readthedocs.io/), a desktop app that guides users through the process of converting data to NWB without writing any code. Using Spyglass could also help with standardization efforts across labs: having a database makes it easy to create lists of names used to refer to particular items and to then move toward standardization.

We also provide many tutorials on the documentation website so that the user can efficiently set up a database and learn to use Spyglass. We continue to actively maintain Spyglass and are eager to work with the community to extend it and support data types and analyses beyond what is currently available. These efforts will increase the usability and reach of Spyglass and make its adoption more attractive, particularly to early-stage investigators. Finally, even in cases where reproducing a result would require installing older versions of software, the results themselves remain accessible within NWB files reference in Spyglass, ensuring that previous results can be built on even as packages evolve.

### Future applications

Spyglass and similar tools have the potential to transform scientific data analysis. In addition to facilitating examination or extension of published results, they enable meta-analysis across studies and easy testing of novel methods across multiple datasets. The machine-readable form of data and analysis pipelines also opens doors for machine-driven analysis and hypothesis testing. As these tools develop and become more accessible, we believe that frameworks like Spyglass will likely become essential for neuroscience researchers.

## Acknowledgements

We thank members of the Frank and Gillespie laboratory for bug reports and testing. We also thank Peter Petersen for consultations about analyzing his publicly available data, Daniel Liu for initial discussions about standardization of pipelines, and Abhilasha Joshi for consulting on the DeepLabCut pipeline. Finally, we thank Vanessa Bender for comments on the manuscript. This work was supported by HHMI funds and NIH grants RF1MH130623 and RF1MH133778 to L.M.F. and a Helen Hay Whitney Postdoctoral Fellowship and NIH grant 1K99EY036953 to K.H.L.

## Author contributions

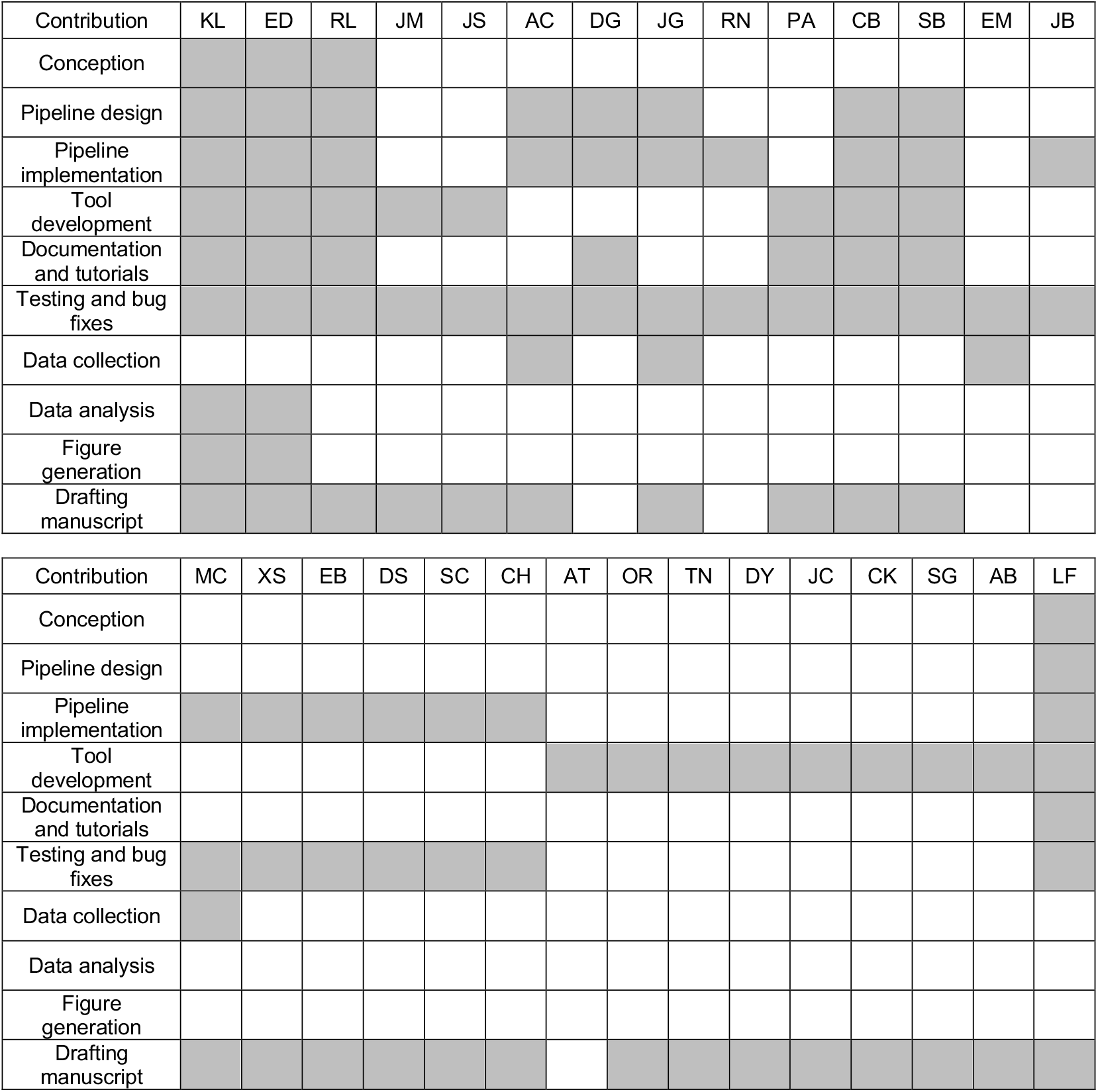

## Methods and materials

### Coding environment

Spyglass was developed in Python 3.9 and is compatible with version 3.10 as well. See our [dependency list](https://github.com/LorenFrankLab/spyglass/network/dependencies) for a full list of Python packages used.

### NWB conversion

To facilitate conversion of raw data to NWB format, we offer [trodes-to-NWB](https://github.com/LorenFrankLab/trodes_to_nwb), a sister package to Spyglass for converting data acquired with the SpikeGadgets hardware to NWB. This comes with a web-based GUI for conveniently generating a YAML file containing the metadata used by `trodes-to-NWB`. For converting data not acquired with SpikeGadgets, users can use NWB conversion tools developed by the NWB team, such as [NeuroConv](https://neuroconv.readthedocs.io/) and [NWB GUIDE](https://nwb-guide.readthedocs.io/).

### NWB file conventions

We adopted a specific set of conventions for our NWB files. Some of these conventions rely on a specific set of [Frank lab-specific NWB extensions](https://github.com/LorenFrankLab/ndx-franklab-novela):

- Time:
  - Spyglass inherits from the source NWB file either the explicit or implicit timestamps. NWB files from Frank lab have explicit timestamps for each sample in Unix time (seconds since 12:00 am January 1^st^, 1970). This lets users to know exactly when data were collected. Spyglass is also compatible with other approaches, however, including implicit timestamping consisting of the start time and sampling rate.
- `ElectrodeTable` and `ElectrodeGroup`:
  - `ElectrodeGroups` are stored in a custom NWB extension that also includes the name of the targeted brain region for each group.
  - The NWB file contains information about the relative locations of each of the electrodes within each physical device used for data collection. This ensures that the relative locations of the electrodes are available for spike sorting and registration to histology.
- Video files
  - The relative path to the video files collected along with the recordings are stored in the NWB file.
- Additional files
  - Other files important to recreate the conditions of the experiments can be saved, depending on the format. For example, the code used for implementing the behavioral paradigm or reward contingency can be stored as text objects in the NWB file.

### NWB file ingestion

Although the NWB format serves as a community standard for neurophysiology data and has a list of [best practices](https://www.nwb.org/best-practices/), it allows some flexibility in the specification of data within NWB files to accommodate user preferences. For example, the `ElectricalSeries` object that stores the electrophysiology data may have different names depending on the convention chosen by the investigator, which may complicate programmatic access to the data. To make Spyglass interoperable with NWB files of varying degrees of NWB-compliance, we have created an option to supply or override information that is missing in the NWB file but is nevertheless required by Spyglass via a configuration file that can accompany the NWB file. We provide an example of this approach in a tutorial (Table 1, 02_Insert_Data).

### Permission-handling and cautious delete

Spyglass is based on a relational database that is accessible to multiple users. In some cases, the type of operations that can be applied to individual data entries (i.e., rows of a table) may need to be restricted to a specified set of users. This is particularly true for operations that are irreversible or time consuming, such as deleting a row from a table storing analysis results. However, there is no inherent mechanism within MySQL or DataJoint that allows permission handling at the level of individual rows of a table. To solve this problem, we have implemented a `cautious_delete` function, in which the user’s permission to carry out a delete operation is checked before it is applied. The permission is granted based on team membership within the lab, reflected in the `LabTeam` table. Though this is not a formal permission-management system, it serves to prevent accidental deletions. We note that this system does incur additional overhead, and while that has not been an issue for us, it is possible that this would become problematic in use for much larger cross-laboratory collaborations.

### Sharing files via Kachery

One way to share the results of Spyglass analysis pipelines is to make the database publicly available. This gives anyone the permission to access the rows of the tables that make up the pipelines and inspect the metadata and the parameters associated with each step of the analysis. But because Spyglass only saves a path to the NWB files containing analysis results within the tables, external viewers cannot download the data and examine it by default.

To enable controlled external access to the data, we have created a system to share selected analysis NWB files with a specified group of users via Kachery. We define a set of tables (`KacheryZone` and `AnalysisNWBfileKachery`) where users can associate analysis NWB files to be shared with a Kachery Zone, making it available to all remote clients who are members of the zone through cloud storage services like Cloudflare R2 bucket or self-hosted servers. Once linked, Spyglass automatically requests, downloads, and manages analysis data for remote users attempting to access shared data through Spyglass tables. This provides a convenient way to provide access to the Spyglass pipelines and associated data files to collaborators.

### Customizing pipelines

To alleviate the challenges associated with database design, we have identified design principles that have been tested extensively by multiple users in the Frank lab. These are described in the text and illustrated with examples in Figures 2 and 3. We recommend users adopt these design elements for building their custom pipelines. We also describe the naming conventions for the tables defined as Python classes and important methods associated with them (e.g. for multiple versions of a pipeline) in our [Developer Notes](https://github.com/LorenFrankLab/spyglass/blob/master/docs/src/contribute.md) available online. Once the pipeline is sufficiently mature and potentially useful to other scientists, we encourage users to submit their pipelines as a pull request to our [GitHub repository](https://github.com/LorenFrankLab/spyglass/tree/master).

### Decoding of position from NWB files from multiple laboratories

The Frank lab data is available on the DANDI archive ([DANDI:000937](https://dandiarchive.org/dandiset/000937/draft)). The Buzsáki lab data was also obtained from DANDI ([DANDI:000059/0.230907.2101](https://doi.org/10.48324/dandi.000059/0.230907.2101)). For decoding the Frank lab data, we applied the clusterless decoding pipeline by detecting the amplitude of threshold-crossing events in the tetrode recordings. For decoding the Buzsáki lab data, we applied a sorted-spikes decoding pipeline. The code for these decoding pipelines, as well as detailed tutorials describing them, are available online (Table 1, 40_Extracting_Clusterless_Waveform_Features, 41_Decoding_Clusterless, 42_Decoding_SortedSpikes). Code to generate Figure 5 can be found at: https://github.com/LorenFrankLab/spyglass-paper. Briefly, decoding the latent neural position and extracting the distance between the most likely decoded position and the animal’s position used methods described in Denovellis et al. (2021). We used a timestep of 4 ms and a position bin size of 2 cm with a continuous (6 cm variance Gaussian random walk) and fragmented (uniform distribution) discrete state. Place intensity receptive fields were estimated using a Gaussian kernel density estimate with a standard deviation of 6 cm for position and 24 mV for amplitude space (amplitude space was used for the clusterless analysis only). We calculated the power of the multiunit firing rate and the decoded distance from the animal by using a multitaper spectrogram during the pre-cooling and cooling periods. The time resolution was 3 seconds and the frequency resolution of 2/3 Hz with a single taper. We excluded immobility periods by using a threshold of 10 cm/s. Power difference was calculated by converting to the Decibel scale and taking the difference of average power under the cooling and pre-cooling condition. The decoded speed of theta sequences was calculated by taking the absolute value of the second-order difference of the decoded distance from the animal (function numpy.gradient) multiplied by the sampling frequency (250 Hz).

## Supplemental Figures

**Supplemental Figure 1:**
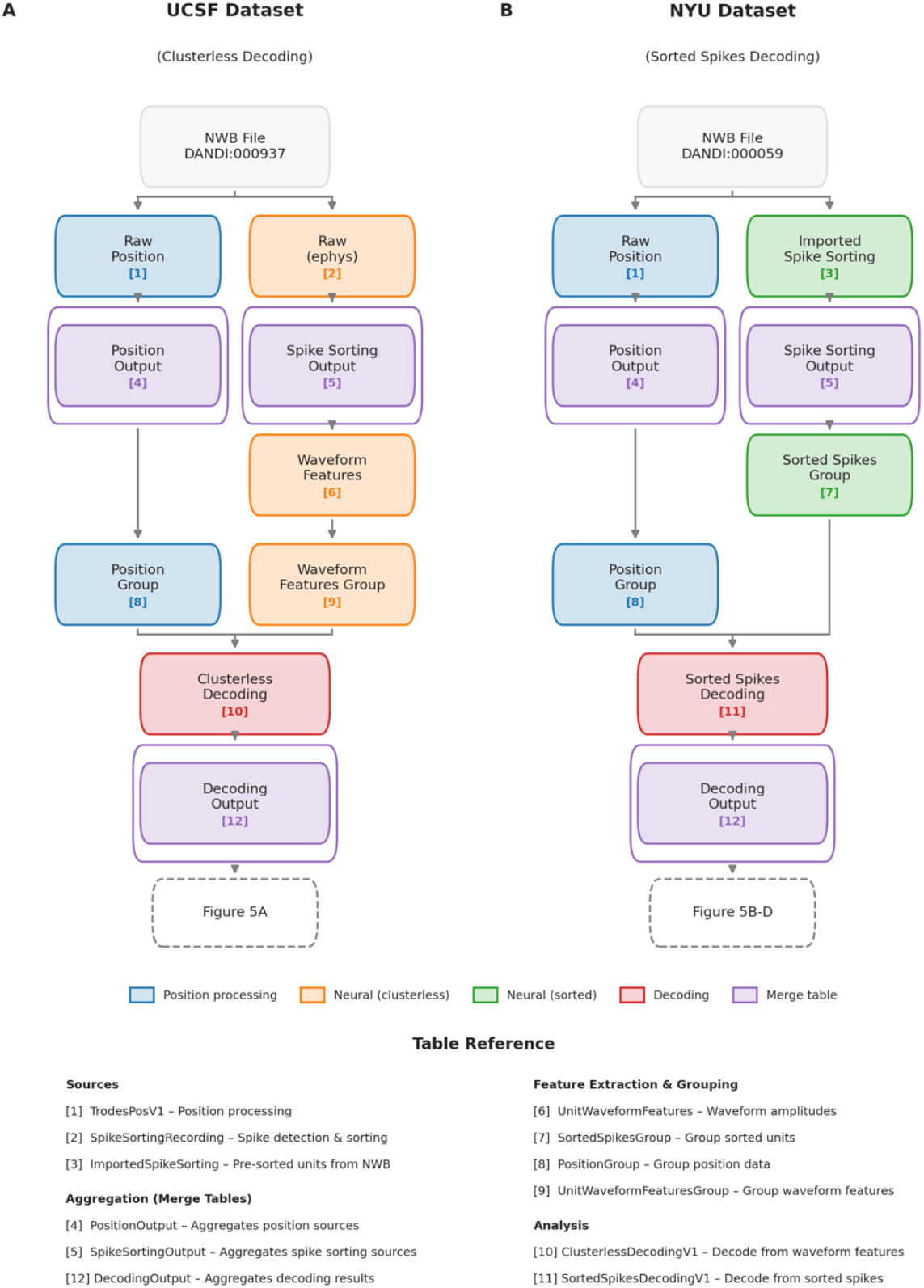
Spyglass pipeline workflow for Figure 5A and B. (A) Decoding of the UCSF dataset started with the NWB file. Data was ingested into the RawPosition and Raw tables, which hold the unprocessed position data (only LED tracking from the Trodes hardware system) and the electrophysiology traces respectively. Position data from the two LEDs had outliers removed, interpolated over, and then smoothed and combined into a single head position via the TrodesPosV1 table (and requiste Selection and Parameter tables which specified the dataset and the parameters for processing). This was then inserted into the PositionOutput merge table. The position data was then inserted into the PositionGroup table which in this case is a passthrough table (but in other cases could hold position data from multiple time periods such as sleep). The raw electrophysiology data was processed through the Spike Sorting pipeline. Because the data is intended for “clusterless” decoding, this simply consists of thresholding for high amplitude spikes (above 60 mV). The UnitWaveformFeatures table then extracts a snippet of waveform data around the time of a spike for each tetrode. UnitWaveformFeature then calculates the peak amplitude at the time of the spike for each tetrode. This amplitude waveform feature (along with the spike time) is used for clusterless decoding in conjunction to the position of the animal via the ClusterlessDecodingV1 table. The decoding result was then ingested into the DecodingOutput merge table which the Figure 5A code subsequently fetched from. (B) The NYU dataset was downloaded from the DANDI archive. The raw position underwent the same processing as Figure 5A. The NYU dataset did not contain raw electrophysiology signals but did contain spike times from already sorted neurons. These were ingested into the ImportedSpikeSorting table and then passed to the SpikeSortingOutput table. The SpikeSortingGroup table allowed us to select only the CA1 cells for decoding. This along with the processed position data was used for decoding via the SortedSpikesDecodingV1 table and inserted into the DecodingOutput merge table. This data was used to generate Figure 5B-D.

